# Identification and Functional Characterization of *Isoflavone Synthase* Gene Family in Pea (*Pisum sativum*): The Entry Point to Pisatin Biosynthesis

**DOI:** 10.64898/2025.12.22.696109

**Authors:** M.S. Tahir, K. Kuflu, N.S. Islam, T. McDowell, S. Dhaubhadel

## Abstract

Isoflavone synthase (IFS), a cytochrome P450 monooxygenase of the CYP93C subfamily, catalyzes the conversion of flavanones into isoflavones, the first committed step in the biosynthesis of isoflavonoid phytoalexins. In pea (*Pisum sativum* L.), the phytoalexin pisatin plays a pivotal role in defense against pathogens. However, the molecular basis underlying IFS function in pea remains poorly understood. In this study, we performed a comprehensive genome-wide identification and characterization of *IFS* genes in pea. Three *IFS* candidates, *PsIFS7A*, *PsIFS7B*, and *PsIFS7C*, were identified that reside on chromosome 7, each harboring all conserved cytochrome P450 signature motifs. *PsIFS* genes exhibited predominant expression in root tissue, with transcript levels induced rapidly upon *Aphanomyces euteiches* infection. Enzymatic assays confirmed their catalytic activity in converting the flavanones naringenin and liquiritigenin into the isoflavones genistein and daidzein, respectively, both *in vitro* and *in planta* systems. Furthermore, all three *PsIFS* genes were found in close proximity to quantitative trait loci (QTL) associated with Aphanomyces root rot resistance. Together, these findings provide novel insights into the *IFS* gene family in pea and lay a foundation for metabolic engineering or molecular breeding strategies to enhance disease resistance through targeted modulation of pisatin biosynthesis.

## 1. Introduction

Pea (*Pisum sativum* L.) is a globally important crop belonging to the Fabaceae family and accounts for approximately 15% of total global pulse production, ranking as the third most cultivated grain legume after common beans and chickpea (Windsor et al., 2024). Since Mendel’s pioneering experiments that laid the foundation of modern genetics, pea has remained a crop of both scientific and agricultural importance. Its balanced nutrient composition, characterized by high levels of protein, starch, and dietary fiber, makes both dry and green peas important in human diets as well as livestock feed (Dahl et al., 2012). As a nitrogen-fixing legume, pea reduces the need for synthetic fertilizers and lowers greenhouse gas emissions, thus supporting sustainable agricultural practices by enhancing soil fertility, facilitating crop rotation, and reducing environmental impact (Bedoussac et al., 2015).

Despite rising global demand, pea yield per unit area remains suboptimal (Trenk et al., 2024). Climate change poses a major challenge in pea cultivation by increasing biotic and abiotic stresses. As a cool-season annual with a relatively shallow root system, pea is particularly vulnerable to root rot disease caused by soil-borne pathogens. Among these, the oomycete *Aphanomyces euteiches* is one of the most destructive pathogens which severely reduces pea production, particularly in Europe and North America (Chatterton et al., 2015; Gaulin et al., 2007; Zitnick-Anderson et al., 2020). *A. euteiches* infects its host plant at any developmental stage, causing the rotting of root and epicotyl, seedling stunting, leaf yellowing, and often plant death (Pfender & Hagedorn, 1982). Under severe Aphanomyces root rot (ARR) pressure during prolonged wet seasons, pea yield can decline by up to 70% (Wu et al., 2018).

Cultural practices, fungicides and biocontrols have shown limited success against ARR due to the soil-borne and persistent nature of the pathogen (Chatterton et al., 2023; Willsey et al., 2021). Consequently, genetic resistance has been identified as the most sustainable strategy (Gangneux et al., 2014; M.-L. Pilet-Nayel et al., 2017). Although complete resistance to *A. euteiches* has not yet been identified in pea, partial resistance has been observed in certain cultivars, and several quantitative trait loci (QTL) linked to ARR resistance have been mapped (Hamon et al., 2011; M.-L. Pilet-Nayel et al., 2005; M. Pilet-Nayel et al., 2002). Elucidating the molecular mechanisms underlying partial resistance is, therefore, essential for guiding breeding programs and developing effective disease management strategies.

Plants produce phytoalexins as part of their defense response against pathogen attack. In legumes, isoflavonoids are the predominant class of specialized metabolites that function as phytoalexins such as medicarpin in alfalfa (Dakora et al., 1993), glyceollin in soybean (Ayers et al., 1976), phaseollin in common bean (Rathmell & Bendall, 1971) and pisatin in pea (Cruickshank & Perrin, 1960). Although pisatin was the first phytoalexin to be chemically characterized (Perrin & Bottomley, 1962), and several radio-labelling studies have sought to elucidate its biosynthesis (Banks & Dewick, 1982; Preisig et al., 1990), previous investigations have only proposed potential precursor compounds involved in pisatin biosynthesis. The order of biosynthetic intermediates and the enzymes catalyzing the key reactions remain largely unresolved.

Isoflavone synthase (IFS), a cytochrome P450 monooxygenase (P450) of the CYP93C subfamily, introduces the isoflavonoid branch into the flavonoid biosynthetic pathway and represents the first committed step in isoflavonoid biosynthesis (**Figure 1**) (Dhaubhadel et al., 2003). In soybean, IFS catalyzes the conversion of naringenin and liquiritigenin into the isoflavone aglycones genistein and daidzein, respectively (Jung et al., 2000; Pandey et al., 2014). These core isoflavones act as the initial precursors for the biosynthesis of more complex isoflavonoids that play critical roles in plant defense (Misra et al., 2010; Samac & Graham, 2007; Subramanian et al., 2007). To date, eight CYP93C subfamily of proteins have been characterized across legume species. In pea, CYP93C18 has been reported to possess IFS activity (Picmanova et al., 2013). Its transcript accumulation and pisatin level increased in response to bruchins, an insect-derived elicitor (Cooper et al., 2005). These findings were made before the availability of the pea whole genome sequence.

**Figure 1.**
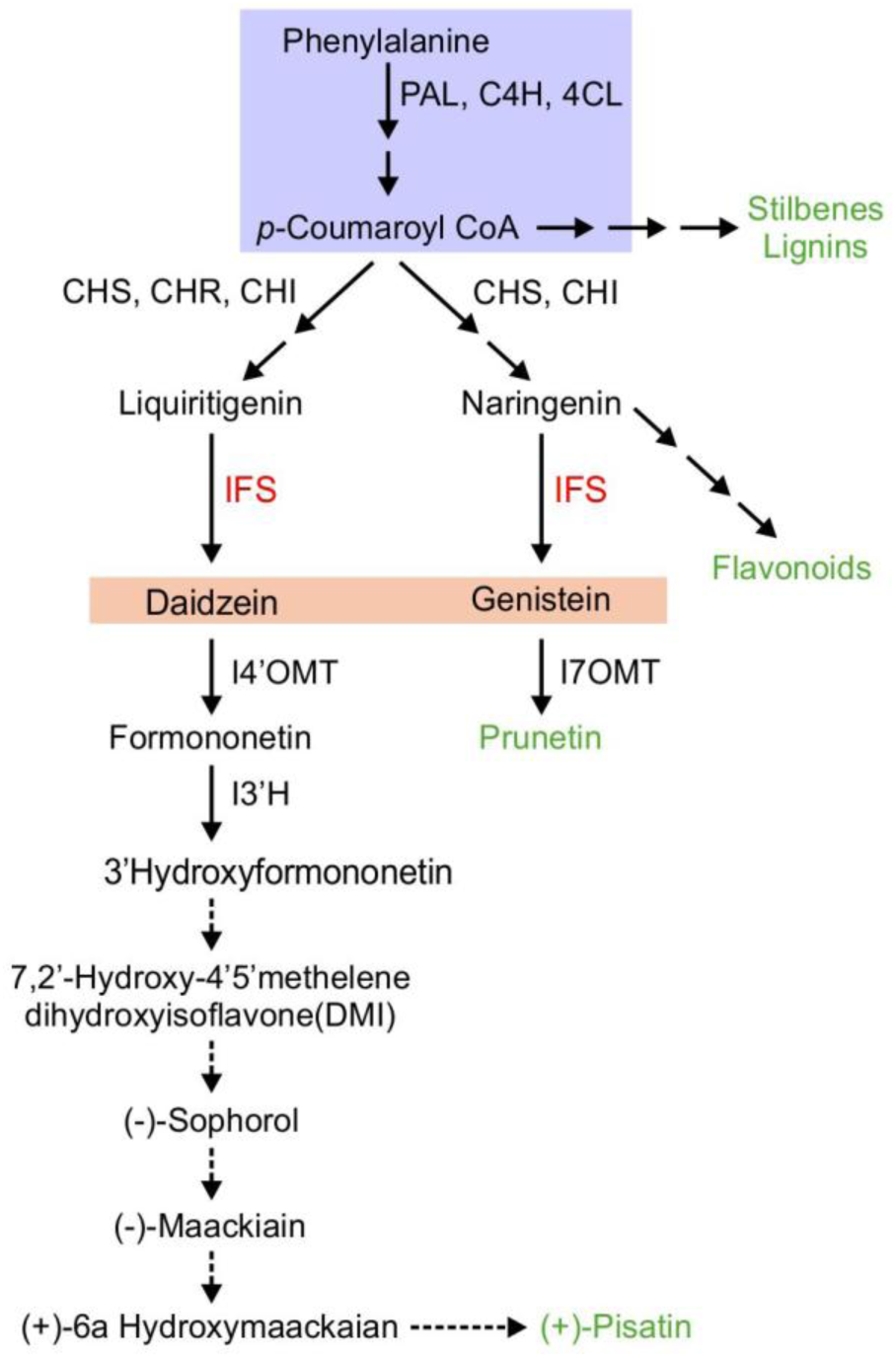
A hypothesized biosynthetic pathway of pisatin in pea. The pathway starts with the general phenylpropanoid pathway (lavender color box) and branches into multiple pathways leading to the production of diverse specialized metabolites (green). The multiple arrows indicate two or more steps in the pathway, and dotted arrows represent speculative steps. The activity of isoflavone synthase (IFS) and other enzymes produces the isoflavones daidzein and genistein. Other enzymes labeled: PAL, phenylalanine ammonia lyase; C4H, cinnamate 4–hydroxylase; 4CL, 4–coumarate:CoA ligase; CHS, chalcone synthase; CHR, chalcone reductase; CHI, chalcone isomerase; I4’OMT, isoflavone 4’-O-methyltransferase; I7OMT, isoflavone 7-O-methyltransferase; I2’H, isoflavone 2’-hydroxylase.

With the release of whole-genome sequences of the two pea cultivars, Caméor (2019) and Zhongwan6 (2023), new opportunities have emerged for genome-wide investigation of genes involved in traits of economic importance. In this study, we leveraged these genomic resources to identify and functionally characterize *IFS* genes in pea (*PsIFS*), with the goal of better understanding of their roles in pisatin biosynthesis and defense. We report that pea harbors three functional *IFS* genes that belong to the CYP93C family. All three *PsIFS* genes are located on chromosome 7, exhibit root-specific expression, and are induced upon pathogen infection. Both *in vitro* enzyme assays and *in planta* studies confirmed their IFS activity. The identification of the functional *PsIFS* genes provides the missing molecular link that initiates isoflavonoid biosynthesis in pea, establishes a foundation for elucidating the complete pisatin pathway and opens avenues for developing enhanced ARR resistance in pea.

## 2 Materials and Methods

### 2.1 Plant materials and *A. euteiches* treatment

Pea (*Pisum sativum* L.) cultivar AAC Chrome seeds were provided by Dr. Syama Chatterton, Lethbridge Research and Development Centre, Agriculture and Agri-Food Canada. Seeds were planted in pots containing medium size sterile vermiculite in a growth chamber with 65% relative humidity at 25°C and 250-400 μmol photons m^-2^s^-1^ light for 16 h and 22°C under dark for 8 h. After two weeks, pea seedlings were transferred to a hydroponic system (Halane et al., 2025), and inoculated with *A. euteiches* zoospores (∼2×10^4^ cells/mL). Mock inoculation contained salt solutions lacking the pathogen. Root samples were collected at 1 and 24 h post-infection, flash frozen in liquid N_2_, and stored at -80°C. Plants were monitored for 14 days post-infection to confirm that the zoospore inoculation resulted in disease symptom.

### 2.2 *In silico* analysis

Since IFS proteins belong to the CYP93C family, we searched the pea (cv Caméor) genome sequences available in the Pulse Crop Database (https://www.pulsedb.org/organism/639) to identify PsIFS candidates. A Basic Local Alignment Search Tool-Protein (BLASTP) against the proteomes of *P. sativum* cv Caméor (Caméor v1a) was performed using the characterized CYP93Cs as query sequences from *Glycine max* (Jung et al., 2000), *Glycyrrhiza echinata* (Akashi et al., 1998), *Glycyrrhiza uralensis* (Li et al., 2024), and *Medicago truncatula* (Chang et al., 2010) (**Supplementary Table S1**). The BLASTP hits with bit score greater than 100 were further analyzed. To refine the search, recurring or duplicate candidates were removed from the dataset, ensuring that only unique and relevant candidates were retained for further analysis. Candidates with more than 40% sequence identity to the query sequences and greater than 400 amino acid residues long were selected for further analysis. The isoelectric point (pI) and molecular mass of selected candidates were calculated using the Expasy online tool, Compute PI/Mw (https://web.expasy.org/compute_pi/). Multiple sequence alignments of candidate proteins were carried out using ClustalW in MEGA 12 (Kumar et al., 2024) and visualized using BOXSHADE 3.21 (https://www.ch.embnet.org/software/BOX_form.html).

To generate a phylogenetic tree, amino acid sequences of CYP93 family of proteins from multiple legumes (**Supplementary Table S2**) were obtained from UniProtKB (https://www.uniprot.org/) or from Du et al. (2016). The tree was generated using the maximum likelihood method with adaptive bootstrapping in MEGA 12 (Kumar et al., 2024).

An AlphaFold2-based Plant Cytochrome P450 Protein Structure Prediction Database (PCPD) was used to predict the protein structures of PsIFS candidates with an imbedded heme cofactor in the structure (Wang et al., 2021). To prepare protein molecules for docking, all the water molecules were deleted and polar hydrogens and Kollman charges were added to the structures using MGLtools 1.5.7 (https://ccsb.scripps.edu/mgltools/). The gridbox of 50 points in each x, y and z directions, was built near the heme cofactor moiety, with a grid spacing of 1 Å. Ligand molecule structures were downloaded from PubChem database (https://pubchem.ncbi.nlm.nih.gov/) and prepared for docking using pymol (https://pymol.org/2/). Docking was performed using Autodock vina (https://vina.scripps.edu/) (Trott & Olson, 2010) and visualized using BIOVIA Discovery Studio Visualizer (https://discover.3ds.com/discovery-studio-visualizerdownload).

For tissue-specific gene expression analysis, publicly available RNA-seq data from the NCBI BioProject database (https://www.ncbi.nlm.nih.gov/bioproject/) was utilized. Specifically, bioproject PRJNA730094 which contains seed (SRR14741927), pod (SRR14741928), flower bud (SRR14741929), flower (SRR14741930), stem (SRR14741931), tendrils (SRR14741932), leaf (SRR14741933) and root (SRR14741934) was used to obtain transcript per million (TPM) reads which were further analyzed using QIAGEN CLC Genomics Workbench (http://www.clcbio.com/products/clc-genomics-workbench/). The candidate gene IDs were used as a search query in the database to obtain the normalized TPM reads. Heatmaps were generated with log_2_ transformed TPM reads using TBtools v1.10056 (Chen et al., 2020).

### 2.3 RNA extraction and RT-qPCR analysis

Total RNA from pea root tissues was extracted using RNeasy Plant Mini kit (Qiagen). On-column DNA digestion was performed using DNase I (Promega). cDNA was synthesized from 1.0 μg of total RNA using the SuperScript^TM^ IV First-Strand Synthesis System (Invitrogen). For qPCR, cDNA was used as a template with SsoFast EvaGreen Supermix (BioRad) and gene-specific primers (**Supplementary Table S3**). The gene expression data was normalized to *PsActin7* (*Psat5g063760.*3). Each experiment included three to four biological replicates, with three technical replicates for each biological replicate. The data were analyzed using CFX Maestro (Bio-Rad).

### 2.4 Gene cloning

The coding regions of *PsIFS* candidates were amplified with high fidelity Platinum® SuperFi DNA Polymerase (Invitrogen) and gene-specific primers (**Supplementary Table S3**). Plasmid constructs were generated using Gateway technology (Invitrogen, USA). Prior to transfer to the destination vector, sequence integrity of the entry clones was verified by sequencing. Expression clones were generated by recombination between the entry clones and the respective Gateway destination vectors using LR clonase reaction mix (Invitrogen, USA). For yeast enzyme assays, the destination vector pESC-Leu2d-LjCPR1-GW (Khatri et al., 2023) was used and recombinant plasmids were transformed to *Saccharomyces cerevisiae* strain BY4742 using Frozen-EZ Yeast Transformation II Kit (Zymo Research, USA). For over expression in pea hairy roots, the entry clone pDONR-PsIFS7C was recombined with the destination vector pK7WG2D (Karimi et al., 2007) and then transformed into *Agrobacterium rhizogenes* strain AR10 by electroporation.

### 2.5 Hairy root generation

For hairy root generation, pea cv AAC Chrome seeds were surface sterilized and grown in sealed magenta boxes containing plant growth medium (2.165 g/L MS salt mixture, 3% sucrose, 0.7% agar pH 5.7) for up to 12 days under 18 h day (22°C), and 8 h night (20°C) cycle.

Explants were inoculated with *A. rhizogenes* AR10 harbouring either the empty vector pK7WG2D or pK7WG2D-PsIFS7C and maintained under the growth condition as described above. After 2-4 weeks post-inoculation, fluorescent transgenic hairy roots were identified under UV-illuminated Nikon SMZ25 stereomicroscope (Nikon Instruments Inc, USA), sub-cultured and maintained on root growth medium (2.165 g/L MS, 3% sucrose, 0.7% agar (w/v), 2.5× MS vitamin mixture, pH 5.7) with 50 μg/mL of Kanamycin (Sigma, USA) and 250 μg/mL of Cefotaxime (GoldBio, USA). For pathogen infection, hairy roots were incubated in mineral salt solution (0.26 g CaCl_2_.2H_2_O/L, 0.07 g KCl/L, 0.49 g MgSO_4_.7H_2_O/L) containing *A. euteiches* zoospores at a concentration of approximately 2×10^4^ cells/mL. After 24 h of infection, roots were flash frozen in liquid N_2_ and stored at -80°C until needed.

### 2.6 Yeast microsome preparation and enzyme assay

For *in vitro* enzyme assay, microsomes were prepared from yeast containing recombinant protein, following the protocols as described previously (Khatri et al., 2023). To examine the catalytic properties of candidate PsIFSs, 1 mg of microsomal proteins were dissolved in 50 mM phosphate buffer (pH 7.6) containing 100 μM naringenin or liquiritigenin substrates (Sigma-Aldrich) and 1 mM NADPH followed by incubation for 6 h at 25°C and 950 rpm. The reaction was stopped by adding equal volume of ice-cold methanol, clarified by centrifugation at 13,000 *g* at 4°C for 5 min and analyzed by HPLC.

To determine the *K*_m_, 20 μg of microsomal protein was incubated with substrate concentrations ranging from 2.5 to 160 μM and 1 mM NADPH at 25°C, 950 rpm for 1 h, followed by HPLC analysis. The reaction products were compared with authentic standards. The *K*_m_ of each candidate PsIFS was estimated from a double reciprocal plot (Lineweaver-Burk plot) of the substrate concentration and the initial velocity of the reaction (1/[S] vs 1/[Vo]).

### 2.7 Metabolite extraction and HPLC/ LC-MS analyses

The enzyme assay reactions were mixed with equal volume of ice-cold methanol and analyzed by HPLC utilizing an Agilent 1260 system, with the following modules binary pump (G7112B), auto-sampler (G7129A), column compartment (G1316A), diode array detector (G4212B). Samples of 10 µL were injected onto a Kinetex XB-C18 (2.6µm x 4.6mm x 100 mm, Phenomenex, USA) maintained at 32°C and peaks were monitored at 280 nm. The mobile phase consisted of solvent A (H_2_O + 0.1% trifluoroacetic acid) and solvent B (Acetonitrile + 0.1% trifluoroacetic acid). Separation occurred at a flow rate of 0.8 mL/min under the following gradient system: 12% B for 1 min, followed by a linear increase of B to 25% over 10 min, increasing to 35% B at 16 min, increasing to 55% B at 17.5 min, increasing to 100% B at 18 min and was held for 1 min. The gradient was then returned to initial conditions over 30 sec and held for 1 min until the next injection for a total analysis time of 20.5 min. The wavelength spectrum and retention time of the substrates and products were compared to commercial standards naringenin, liquiritigenin, genistein, and daidzein (Sigma-Aldrich).

For LC-MS analysis, metabolites were extracted by vortexing lyophilized tissues in 80% methanol (200 µL mg^-1^ tissue) for 30 sec followed by sonication for 15 min. The samples were then incubated in a thermomixer (25°C) at 1200 rpm (Eppendorf ThermoMixer®) for 10 min. The homogenate was centrifuged at 11,000 *g* for 10 min at 4°C and 350 µL of the supernatant was dried under nitrogen gas. Finally, the dried metabolite extracts were redissolved in 50% methanol for analysis.

The samples were run using a Thermo Q-Exactive Orbitrap mass spectrometer coupled to an Agilent 1290 HPLC system (Santa Clara, CA, USA) as described in Sakariyahu et al. (2025). Each sample (2 μL) was injected onto a Zorbax Eclipse Plus RRHD C18 column (2.1 mm × 50 mm, 1.8 μm; Agilent) maintained at 35°C using a flow rate of 0.3 mL min^−1^ with a mobile phase of LC-MS-grade water (Optima) with 0.1% formic acid (phase A) and LC-MS-grade acetonitrile (Optima) with 0.1% formic acid (phase B). Mobile phase B was held at 2% for 45 sec before increasing to 22% over 30 sec. Mobile phase B was increased to 35% over 1.75 min and to 100% over 3.5 min. Mobile phase B was maintained at 100% for 2.5 min before returning to 2% over 30 sec.

Samples were analyzed by a data dependent acquisition (DDA) method that consisted of a full scan at 35,000 resolutions across a 100-1250 *m/z* range, a max injection time (max IT) of 128 ms and an automatic gain control (AGC) of 1 × 10^6^. The top 5 most intense ions of each full MS scan (dynamic exclusion: 10 s) were selected for MS/MS scans at 17,500 resolutions using normalized collision energy at 35 NCE, a maxIT of 64 ms, an AGC of 5 × 10^6^, and an isolation window of 1.2 *m/z* units.

## 3 Results

### 3.1 Pea genome contains three *PsIFS* genes

BLASTP searches of the pea (cv Caméor) proteome using five characterized CYP93Cs identified 824 hits, which were reduced to 154 unique sequences. After excluding proteins with <400 amino acids and with <40% identity to the query sequences, eight PsIFS candidates were selected for further analysis (**Supplementary Table S1, Supplementary Figure S1)**.

To validate if all eight PsIFS candidates belong to CYP93C family, a phylogenetic tree was constructed using the eight putative PsIFSs along with other CYP93 protein sequences from multiple legume species (**Supplementary Table S2**). As shown in **Figure 2**, the tree branched into three major clades: CYP93B and CYP93C subfamilies grouped into one and CYP93A and CYB93E subfamilies into another clad, while CYP93G formed a distinct group. Three PsIFS candidates Psat7g220280.1, Psat3g159880.1, and Psat7g220160.1 clustered with CYP93B proteins, suggesting their possible function as flavone synthase II enzymes (Fliegmann et al., 2010) while Psat3g070080.1 grouped with CYP93A proteins. Two CYP93As from soybean have been reported to exhibit 3,9-dihydroxypterocarpan 6A-monooxygenase activity (Xia et al., 2023), suggesting a similar function for Psat3g070080.1. Interestingly, Psat4g121920.1 did not cluster with any defined CYP93 subfamily, indicating a possible divergence or unrelated function.

**Figure 2.**
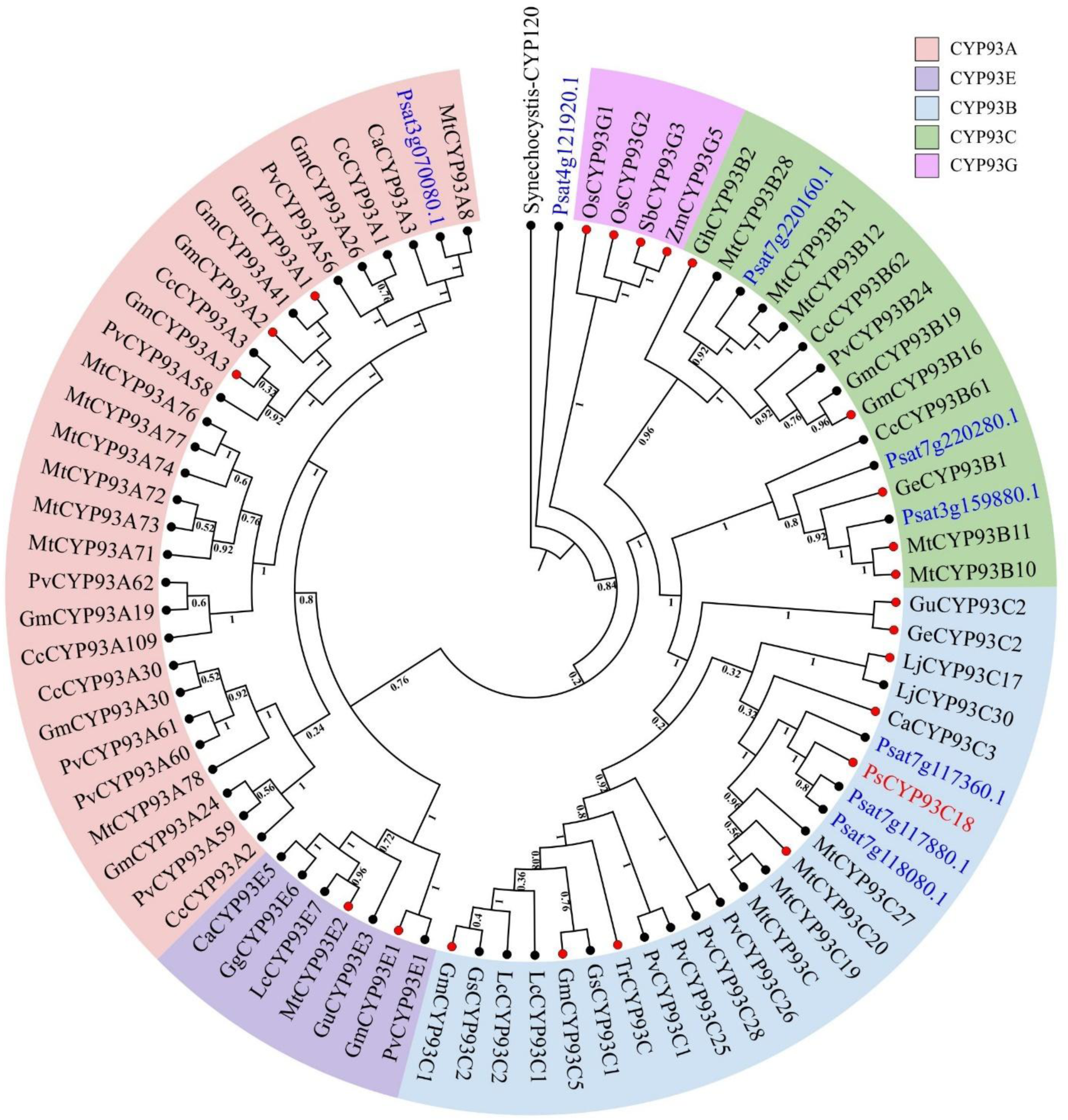
Phylogenetic analysis of CYP93 family proteins and PsIFS candidates. A maximum-likelihood phylogenetic tree was constructed using the deduced protein sequences of the eight PsIFS candidates together with 72 CYP93 family proteins from other plant species. The tree was generated using adaptive bootstrapping and the bootstrap support values are shown next to the branches at a scale of 1.0. Colors indicate CYP93 subfamily as labeled. Functionally characterized CYP93 proteins are marked with red circles at the branch tips, pea candidate proteins are shown in blue font. CYP120 from *Synechocystis* (accession no. 2VE3) was used as an outgroup. Mt, *Medicago truncatula*; Pv, *Phaseolus vulgaris*; Tr, *Trifolium repens*; Gs, *Glycine soja*; Gm, *Glycine max*; Lc, *Lens culinaris*; Gu, *Glycyrrhiza uralensis*; Gg, *Glycyrrhiza glabra*; Ca, *Cicer arietinum*; Cc, *Cajanus cajan*; Os, *Oriza sativa*; Sb, *Sorghum bicolor*; Zm, *Zea mays*; Gh, *Gerbera hybrida*; Ge, *Glycyrrhiza echinata*; Lj, *Lotus japonicus*

CYP93C subfamily of proteins have been shown to possess IFS activity in soybean (Jung et al., 2000; Steele et al., 1999), *Lotus japonicus* (Shimada et al., 2000), *Trifolium repenes* (Franzmayr et al., 2012), and *Glycyrrhiza echinata* (Akashi et al., 1998). Three putative PsIFSs, Psat7g117360 (PsIFS7A), Psat7g117880 (PsIFS7B), and Psat7g118080 (PsIFS7C), clustered with previously characterized CYP93C proteins from pea (CYP93C18) and other plant species suggesting their potential role as IFS in pea. Interestingly, the genomic location of *CYP93C18* could not be found in the whole genome sequence of both pea cultivars Caméor and Zhongwan6. A BLASTP search using CYP93C18 as a query provided only three hits PsIFS7A, PsIFS7B and PsIFS7C with 92.18%, 97.71% and 97.32% amino acid identity, respectively, suggesting that the observed sequence differences could be cultivar-specific polymorphisms. Genes encoding the three PsIFSs are located on chromosome 7 at three different loci. The identification of the putative *PsIFS* gene family and their detailed characteristics are summarized in **Supplementary Figure S1** and **Table 1**, respectively.

**Table 1.** Detailed comparison of *PsIFS* candidates in pea.

| Cultivar | Given name | Gene ID | Gene location | CDS length | Protein length | Protien MW (kDA) | pI |
| --- | --- | --- | --- | --- | --- | --- | --- |
| PeaZW6<sup>2</sup> | *PsIFS7A* | Psat07G0304000 | chr7:201999498..202001223 | 1644 | 547 | 61.97 | 9.02 |
|  | *PsIFS7B* | Psat07G0303200 | chr7:200698898..200700559 | 1575 | 524 | 59.31 | 8.96 |
|  | *PsIFS7C* | Psat07G0303400 | chr7:201019085..201020746 | 1575 | 524 | 59.28 | 8.96 |
| PeaCaméor<sup>1</sup> | *PsIFS7A* | Psat7g117360 | chr7LG7:192487626..192489332 | 1578 | 525 | 59.47 | 8.71 |
|  | *PsIFS7B* | Psat7g117880 | chr7LG7:193479794..193481583 | 1641 | 546 | 61.77 | 9.23 |
|  | *PsIFS7C* | Psat7g118080 | chr7LG7:193800007..193802431 | 1575 | 524 | 59.28 | 8.96 |

<sup>1</sup>pea cv Caméor, <sup>2</sup>pea cv Zhongwan(ZW)6

### 3.2 Sequence analysis of putative *PsIFS*

Sequence analysis of the putative *PsIFS* genes revealed a conserved structure of two exons and one intron, encoding proteins with predicted molecular masses of 59.28 to 61.77 kDa. *PsIFS7C* encodes a protein with 524 amino acid residues that is identical across the two available pea reference cultivars Caméor and Zhongwan6, with a predicted molecular weight of 59.28 kDa (**Table 1, Supplementary Figure S2**). In contrast, PsIFS7A and PsIFS7B displayed notable varietal differences between the reference genomes. The predicted isoelectric points (pI) of PsIFS7A, PsIFS7B, and PsIFS7C ranged from 8.71 to 9.23, consistent with alkaline proteins. All three putative PsIFSs, PsIFS7A, PsIFS7B, and PsIFS7C, are predicted to be localized in the endoplasmic reticulum with high confidence probabilities: 0.92, 0.88, and 0.91, respectively, consistent with their expected role as P450 enzymes (Dastmalchi et al., 2016).

Amino acid sequence alignment of PsIFS candidates from pea cv Caméor with four functionally characterized CYP93Cs from other legumes revealed that the PsIFS proteins contain all conserved motifs of the cytochrome P450 family, with some variations. The third amino acid residue in the proline-rich motif (PPGP) at the N-terminal region exhibits substitution of Gly with Ser in most of the characterized CYP93C subfamily, including PsIFS7A whereas in PsIFS7B and PsIFS7C, Gly is substituted by a Cys residue. The I-helix/oxygen-binding motif (AGTDST), K-helix motif (EXXR), PERF motif, and the C-terminus heme binding site (FGSGRRMCPG) that are signature conserved motifs of P450 subfamilies are conserved in the three PsIFS proteins **(Figure 3**, **Table 2**). Notably, PsIFS7B and PsIFS7C exhibited 100% sequence identity within the five conserved motifs, suggesting functional redundancy or close evolutionary conservation between these two isoforms. PsIFS7B contained a 22 amino acid long repeat sequence at the N terminal end which is missing in PsIFS7A and PsIFS7C candidates and the characterized CYP93C proteins from other legume species. Additionally, pea cv Zhongwan6 reference genome also lacks the 22 amino acid repeat sequence in PsIFS7B (**Supplementary Figure S2**).

**Figure 3.**
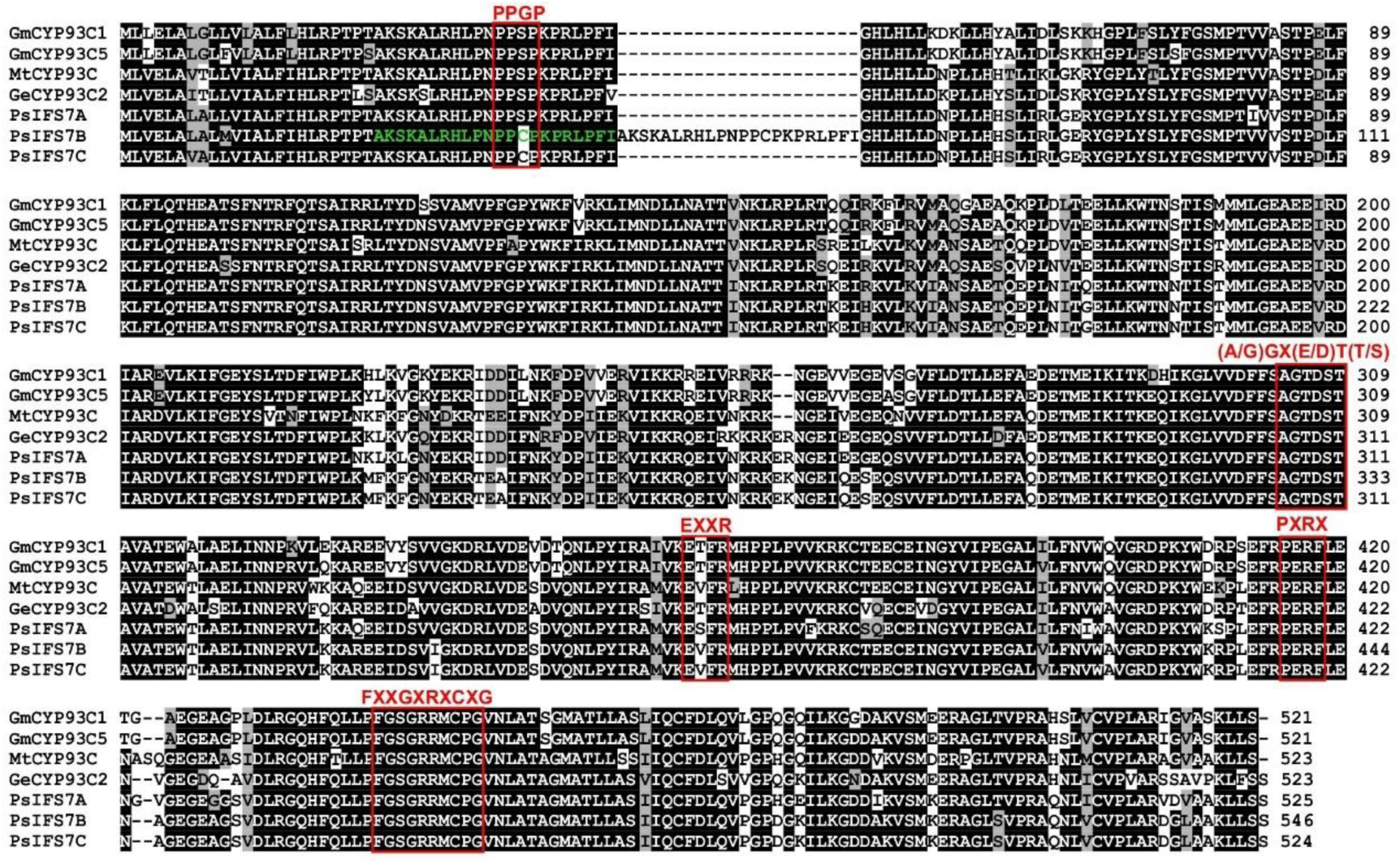
Multiple sequence alignment of PsIFS candidates with functionally characterized CYP93C proteins using ClustalO. Identical and similar amino acid residues are shown in black and gray shading, respectively. Conserved P450 signature motifs are indicated by red boxes. Repeat sequence within PsIFS7B are indicated. Accession numbers of sequences used are: GmCYP93C1 (B5L5C9), GmCYP93C5 (Q9SWR5), MtCYP93C (AY167424), GeCYP93C2 (Q9SXS3), PsIFS7A (Psat7g117360), PsIFS7B (Psat7g117880), and PsIFS7C (Psat7g118080).

**Table 2.**
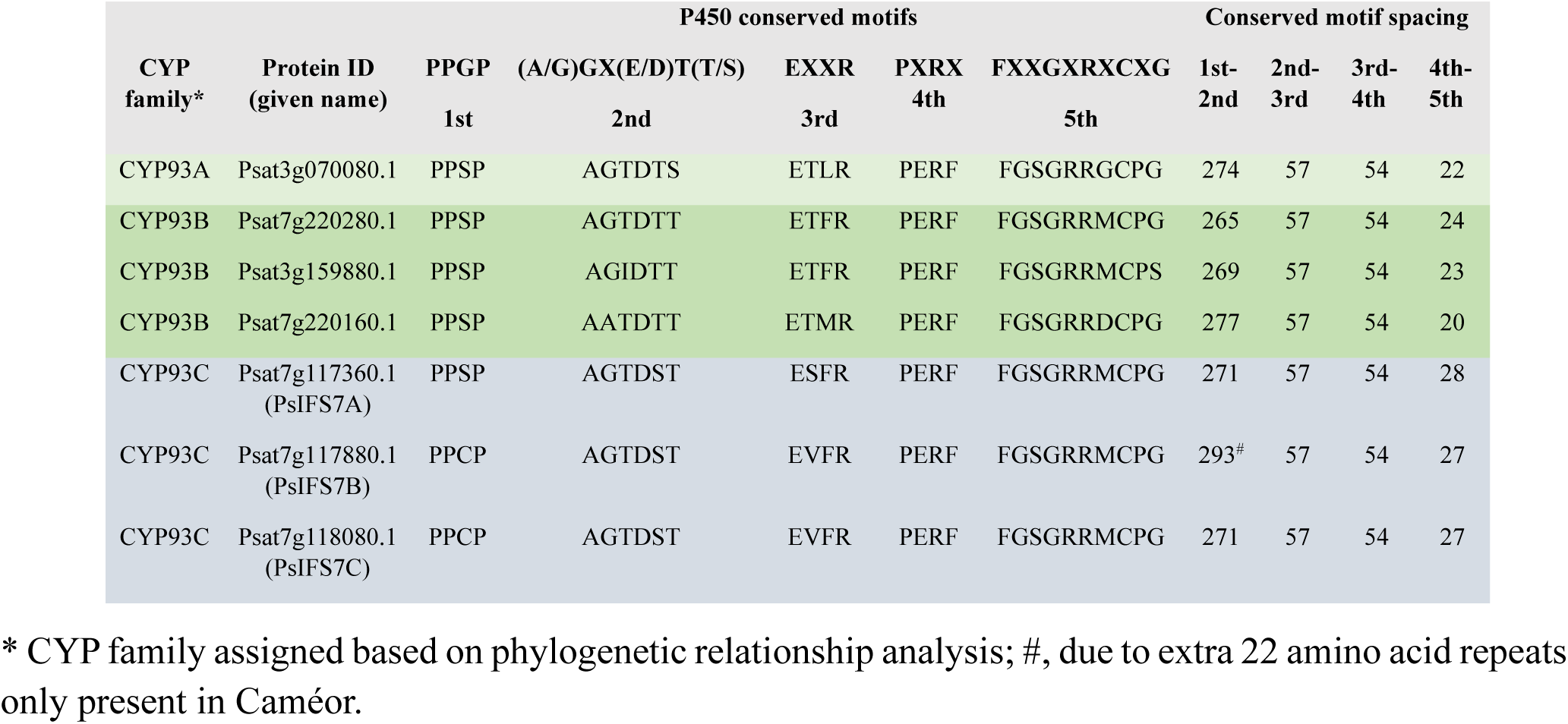
P450 conserved motif analysis of PsIFS candidates in pea cv Caméor.

### 3.3 Molecular modeling of PsIFS candidates identifies substrate-specific interactions

To elucidate the structural basis of enzyme-substrate specificity, homology modeling was performed to predict the 3D structures of PsIFS candidate proteins, followed by molecular docking with naringenin and liquiritigenin as substrates. The analysis revealed strong binding affinities of both substrates to PsIFS7A, PsIFS7B, and PsIFS7C, with estimated binding free energies of –8.26, –7.90, and –7.20 kcal/mol for naringenin, and –7.99, –8.19, and –7.76 kcal/mol for liquiritigenin, respectively. In all three enzymes, the substrates exhibited characteristic Pi-Pi T-shaped interactions between the benzene rings of the flavanone and the heme group, positioning the aromatic ring close to the heme iron in the catalytic pocket (**Figure 4**). In PsIFS7A, Leu371 and Leu500 mediated interactions with both naringenin and liquiritigenin, while Ala120 and Ala306 additionally contributed to binding with liquiritigenin. In PsIFS7B, Ile110, Ala306, Leu371, and Leu499 participated in hydrophobic interactions with both substrates. In PsIFS7C, naringenin interacted with Ile110, Leu371, and Leu499, whereas liquiritigenin formed interactions with Ile110, Ala120, and Ala306. Notably, the critical residue Ala306, located within the oxygen-binding motif of all three PsIFS enzymes, formed Pi-alkyl interactions with liquiritigenin, indicating its key role in substrate binding and catalysis. However, for naringenin, this interaction with Ala306 was observed only in PsIFS7B. Neither PsIFS7A nor PsIFS7B utilized residues from any of the conserved cytochrome P450 signature motifs for interaction with naringenin. Psat7g220280 (PsCYP93E), which did not show favorable substrate orientation towards the heme iron, was used as a negative control, resulting in no Pi-interactions with the substrates.

**Figure 4.**
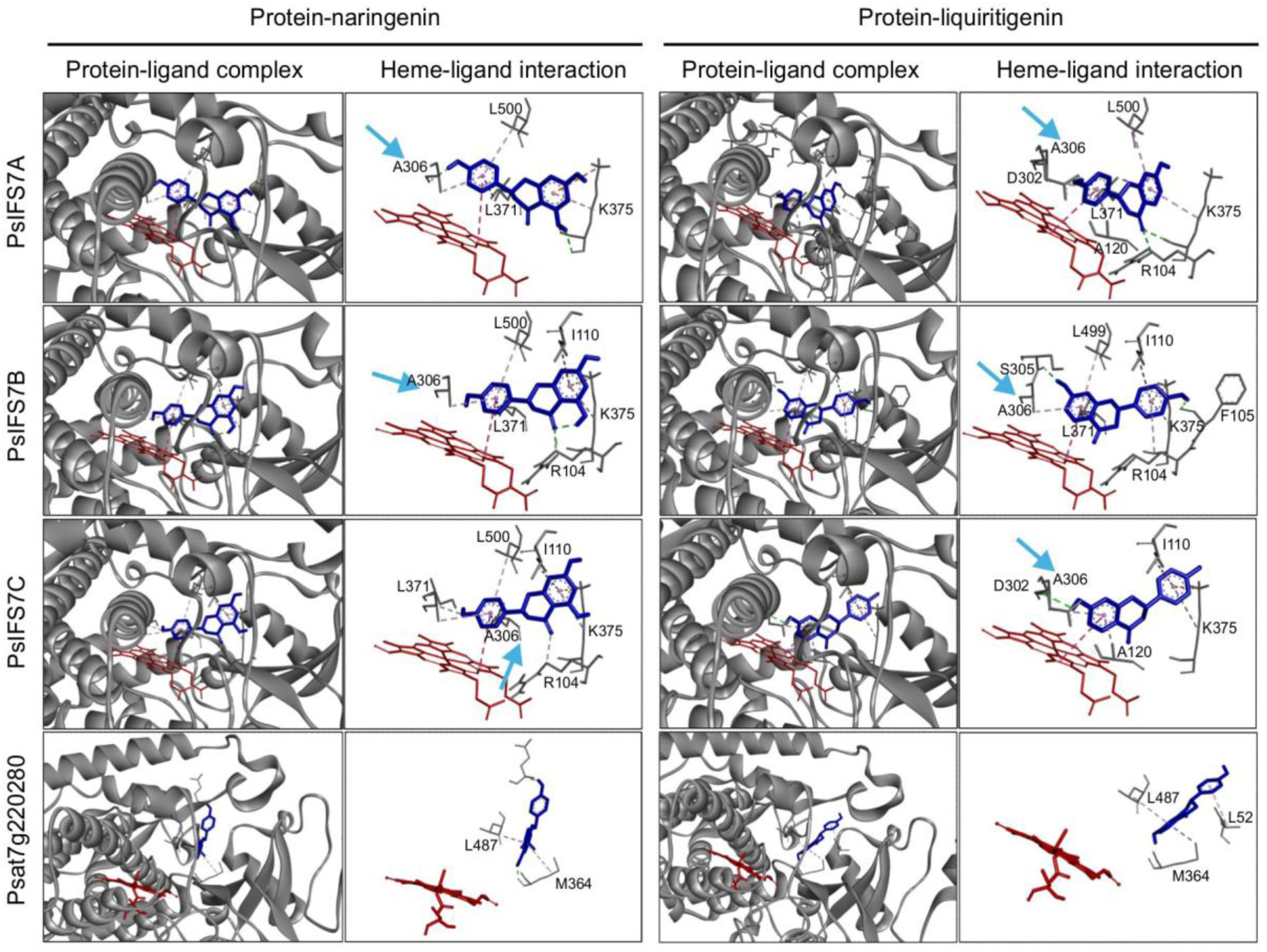
Molecular docking of PsIFSs with naringenin and liquiritigenin as substrates. Predicted three-dimensional structures of PsIFS7A, PsIFS7B, and PsIFS7C proteins docked with their substrates are shown. The overall conformation of the protein-ligand complex, highlighting the substrate (blue) and the heme group (red) within the catalytic pocket is shown. A close-up of hydrophobic and Pi interactions between the substrate and key amino acid residues and heme-group is indicated in heme-ligand interaction. Conserved alanine residue in PsIFSs are indicated by blue arrow. Psat7g220280 (CYP93B-like) was used as a negative control and showed no substrate orientation toward the heme iron.

### 3.4 *PsIFSs* show root-specific gene expression and are induced upon pathogen infection

Since ARR is a soil borne disease with roots exposed to the pathogen first, we assessed the tissue-specific expression of the putative *PsIFS* genes using the publicly available transcriptomic data (BioProject: PRJNA730094) that contains eight pea tissue types including root, leaf, tendril, stem, flower, flower bud, green pod, and immature seed. The analysis revealed that the expression levels of *PsIFS7A*, *PsIFS7B,* and *PsIFS7C* varied across different tissue types with maximum expression found in roots (**Figure 5a)**.

**Figure 5.**
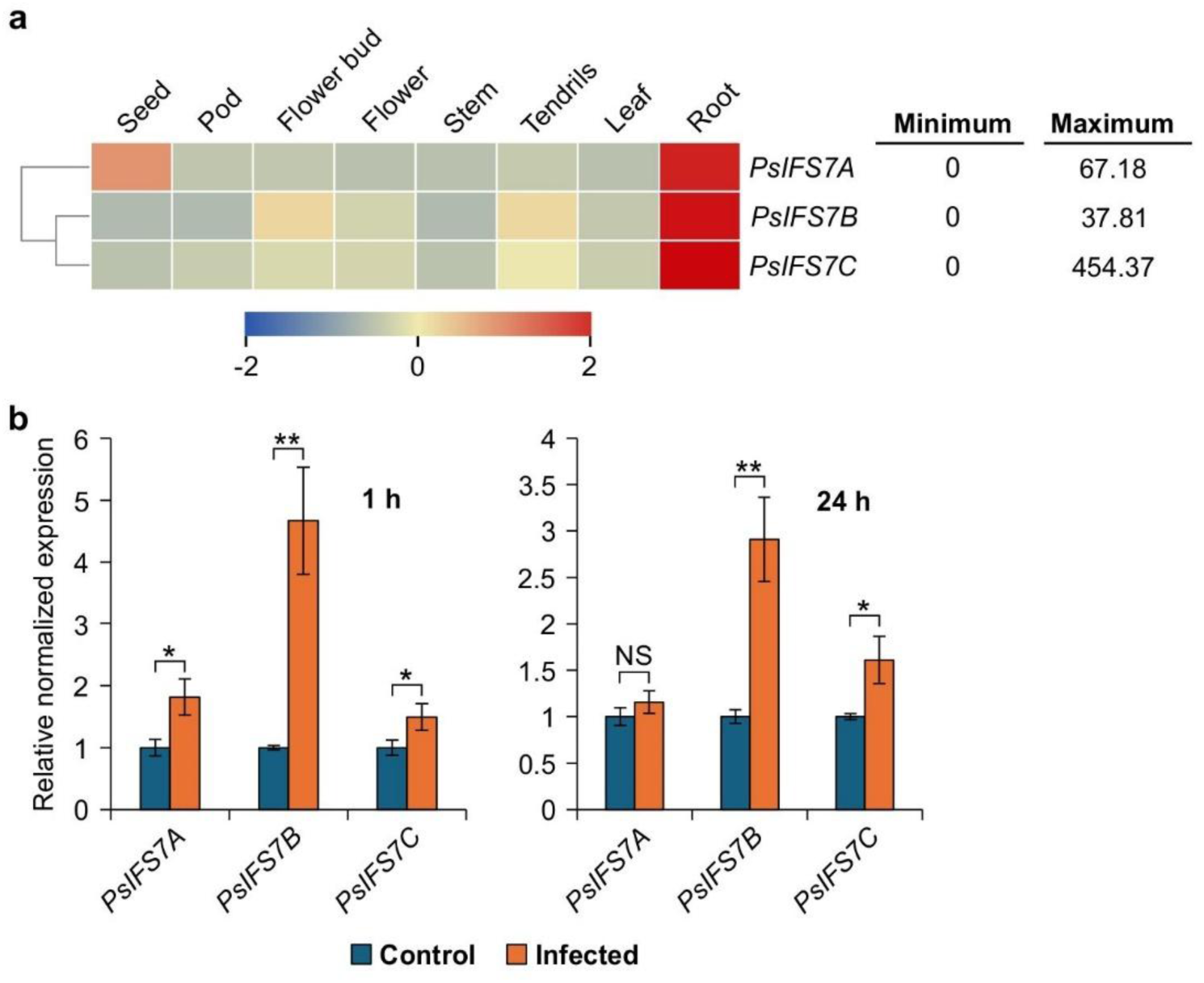
Expression profiling of *PsIFS* genes. (a) Tissue-specific expression of *PsIFS* candidate genes were obtained from the bioproject PRJNA730094 for the heatmap generation. The color scale represents expression values, with blue and red indicating low and high transcript abundance, respectively. Transcripts per million (TPM) values for each gene are shown to the right. The heatmap was generated using log₂-transformed TPM values, normalized across rows. (b) Relative transcript accumulation of *PsIFS* genes in pea roots infected with *A. euteiches*. cDNA was synthesized from total RNA (1 μg) isolated from pea roots post-infection and qPCR was performed using gene-specific primers. Expression values were normalized against the reference gene *PsActin7*. Error bars represent the SEM of four biological replicates, each with three technical replicates. Asterisks indicate significant differences between control and infected samples (\**p*< 0.05; \*\**p*< 0.01).

To investigate the transcriptional response of putative *PsIFS* genes to pathogen infection, pea cv AAC Chrome plants were inoculated with *A. euteiches*, and transcript abundance of *PsIFS* candidates was quantified using qRT-PCR. Mock-inoculated root tissues were included as controls.

The transcript levels of *PsIFS7A*, *PsIFS7B*, and *PsIFS7C* increased significantly at 1 h post-infection in infected roots relative to controls. Transcript accumulation of *PsIFS7B* and *PsIFS7C* remained elevated at 24 h post-infection, whereas *PsIFS7A* expression returned to basal levels (**Figure 5b**). Notably, compared with mock-inoculated roots*, PsIFS7B* transcripts increased 4.6- and 2.9-fold at 1 and 24 h post-infection, respectively. The rapid and sustained induction of *PsIFS7B* and *PsIFS7C* suggests their potential involvement in defense-associated isoflavonoid and phytoalexin biosynthesis during *A. euteiches* infection.

### 3.5 Functional characterization of PsIFSs

To determine the enzymatic function of candidate PsIFSs, *PsIFS7A*, *PsIFS7B*, and *PsIFS7C* genes were cloned individually in a yeast expression vector. Yeast microsomal fractions containing recombinant PsIFS were isolated and used in an *in vitro* enzyme assay in the presence of NADPH to evaluate their ability for hydroxylation and aryl ring migration, thereby producing daidzein and genistein from the substrates liquiritigenin and naringenin, respectively. Product formation and substrate depletion was measured by HPLC analysis using authentic standards. Reactions with microsomes isolated from yeast strain containing an empty vector (vector only), no substrate and no protein were used as the negative controls. Our results demonstrated that all three PsIFSs use liquiritigenin as a substrate to produce daidzein while only PsIFS7B and PsIFS7C were able to catalyze the conversion of naringenin to genistein. No reaction product was formed when microsomes from yeast expressing PsIFS7A were incubated with naringenin (**Figure 6**). Similarly, no reaction products were detected in the negative controls. For the recombinant PsIFS proteins, K_m_ estimates ranged from 5.14 µM (PsIFS7B) to 7.66 µM (PsIFS7C) for naringenin. While K_m_ estimates with liquiritigenin as a substrate were 15.6, 7.0, and 10.7 µM for PsIFS7A, PsIFS7B and PsIFS7C, respectively, suggesting that PsIFS7B had the highest affinity for both the substrates used in the study. We calculated *k_cat_* and catalytic efficiency for each enzyme with both substrates, assuming that the enzyme concentrations in the yeast microsomal protein preparations were identical. The detailed kinetic properties of the PsIFSs are indicated in **Table 3**.

**Figure 6.**
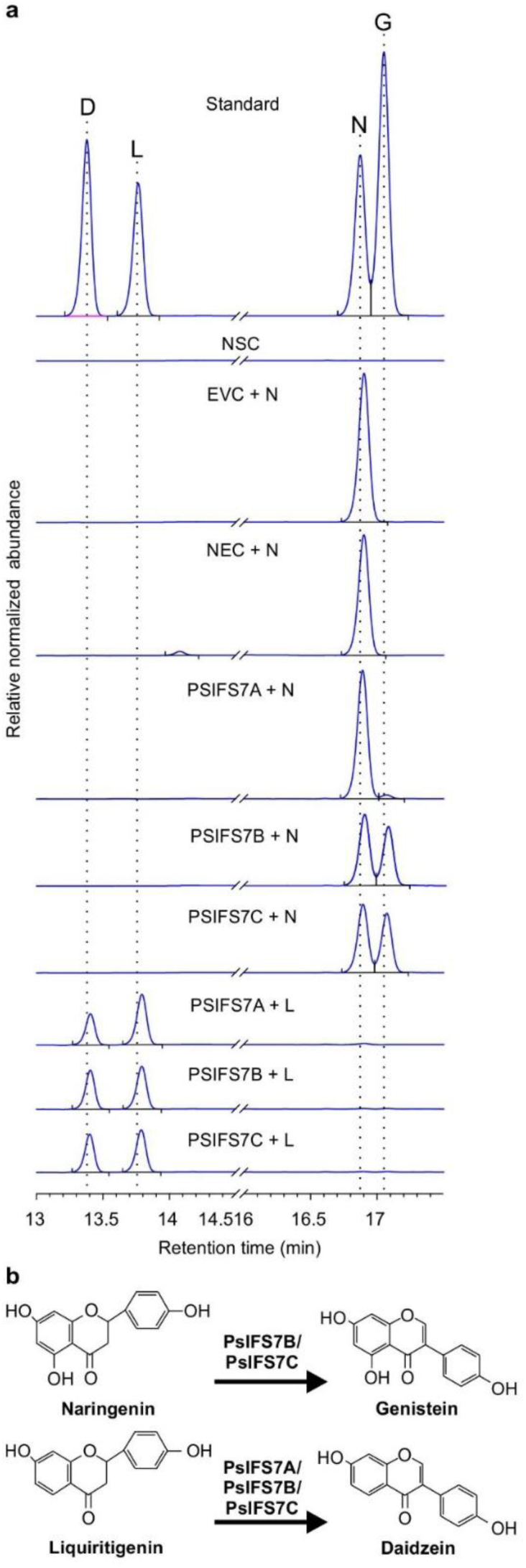
*In vitro* functional characterization of candidate PsIFSs. (a) Yeast microsomal fraction containing candidate PsIFSs and LjCPR1 was incubated with the substrate naringenin or liquiritigenin in the presence of NADPH followed by high-performance liquid chromatography (HPLC) analysis. Standards: D, daidzein; G, genistein; N, naringenin; L, liquiritigenin. NSC, no substrate control; EVC, empty vector control; NEC, no enzyme control. (b) Enzymatic reaction conversion of liquiritigenin to daidzein and naringenin to genistein.

**Table 3.** Kinetic parameters of PsIFS enzymes with flavanone substrates.

| Protein | Substrates | $V_{max}$ (nmol<br>$\text{min}^{-1} \text{mg}^{-1}$) | $K_m$ ($\mu\text{M}$) | $k_{cat}$ ($\text{s}^{-1}$) | Catalytic<br>efficiency<br>($\mu\text{M}^{-1} \text{min}^{-1}$) |
| --- | --- | --- | --- | --- | --- |
| PsIFS7A | Liquiritigenin | $0.466 \pm 0.0184$ | $15.6 \pm 1.08$ | $0.0231 \pm 0.0009$ | $0.0896 \pm 0.0054$ |
|  | Naringenin | nd | nd | nd | nd |
| PsIFS7B | Liquiritigenin | $0.997 \pm 0.0144$ | $7.00 \pm 0.272$ | $0.0493 \pm 0.0007$ | $0.406 \pm 0.0051$ |
| | Naringenin | $0.253 \pm 0.0023$ | $5.14 \pm 0.236$ | $0.0125 \pm 0.0001$ | $0.147 \pm 0.0066$ |
| PsIFS7C | Liquiritigenin | $1.55 \pm 0.011$ | $10.7 \pm 0.151$ | $0.0767 \pm 0.0005$ | $0.428 \pm 0.0053$ |
| | Naringenin | $0.465 \pm 0.0037$ | $7.66 \pm 0.017$ | $0.023 \pm 0.0001$ | $0.18 \pm 0.0016$ |

nd, not determined; $V_{max}$ , $K_m$ and $k_{cat}$ are expressed as mean values of three independent replicates ± SE

Since PsIFS7B and PsIFS7C share 99.62% sequence identity at the amino acid level, to confirm the role of these enzymes *in planta*, we overexpressed *PsIFS7C* using hairy root system in pea. Transgenic hairy roots expressing GFP were selected under a microscope and transcript level of *PsIFS7C* was verified by qPCR (**Figure 7a-c**). Four independent transgenic roots overexpressing *PsIFS7C* (PsIFS7C-OE) or empty vector only (control) were selected for the metabolite analysis (**Figure 7c**). We observed that daidzein level was increased in all PsIFS7C-OE roots compared to the controls (**Figure 7d**). Some discrepancy was observed for PsIFS7C-OE line 4 and control line 1 where genistein accumulation level did not show similar trend as the other PsIFS7C-OE or control lines (**Figure 7e**). Nevertheless, the majority of transgenic PsIFS7C-OE roots contained higher levels of genistein compared to controls. These results confirm that PsIFS7C possesses IFS activity initiating the isoflavonoid branch in the phenylpropanoid pathway in pea.

**Figure 7.**
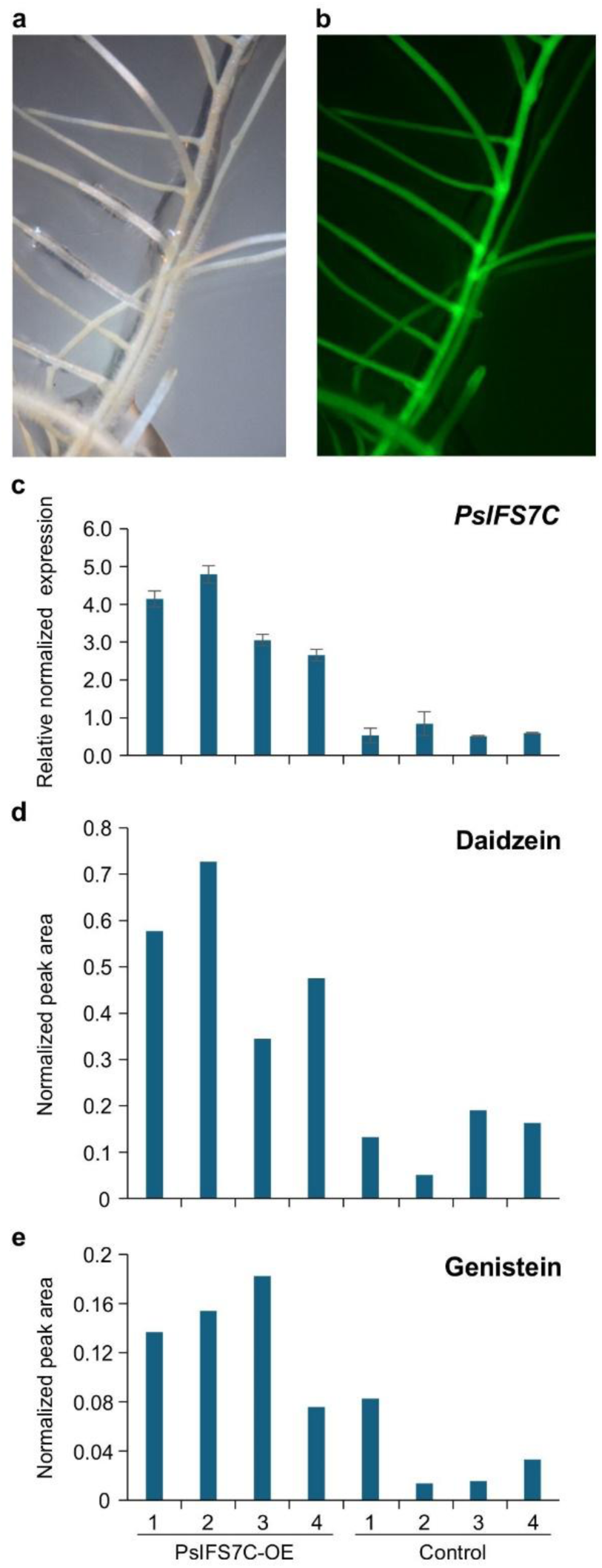
Functional characterization of *PsIFH7C in planta.* Photographs of pea hairy roots under (a) white light and (b) UV light with a GFP filter indicating transgenic roots are shown. (c) *PsIFS7C* transcript levels in four independent PsIFS7C-OE and control hairy roots analyzed using qRT-PCR. (d) Daidzein and (e) Genistein levels in the four PsIFS7C-OE and control hairy roots as measured by LCMS analysis.

To determine whether *PsIFS* genes are linked to QTL associated with ARR resistance, we examined QTL for the trait located on chromosome 7 (∼549 Mbp). The analysis identified five QTL distributed across a ∼154 Mbp region, where several QTL and markers were found to overlap the same genomic location (**Table 4**, **Figure 8**). *PsIFS7A*, *PsIFS7B*, and *PsIFS7C* are clustered within a 1 Mbp region located within 40 Mbp region from the identified QTL on either side. Although these genes do not reside directly within the QTL interval, their close physical proximity suggests a potential genetic linkage between the *PsIFS* cluster and the ARR-associated QTL.

**Table 4.** QTL associated with ARR resistance on Chromosome 7 in pea.

| QTL | Parents | Peak (cM) | Marker Name | Marker genomic location (in ZW6) | Type of Marker | LOD | References |
| --- | --- | --- | --- | --- | --- | --- | --- |
| AeMRCD1Ps-7.1 | Reward'×'00-2067 | 124.1 | PsCam058653_39030_117 (Left) | 142,690,410 | SNP | 5.7 | Wu et. al., 2021 |
|  |  |  | PsCam060132_40119_811 (Right) | 161,608,643 |  |  |  |
| AeMRCD1Ps-7.2 | Reward'×'00-2067 | 211.8 | PsCam033614_19198_1651 (Left) | 242,422,915 | SNP | 5.2 |  |
|  |  |  | PsCam024843_14133_904 (Right) | 295,998,769 |  |  |  |
| Ae-Ps7.5 | Baccara × PI 180693 and Baccara × 552 RIL | 179.5 | AA317 | 253,417,369 | SSR | 4.9 | Hamon et. al., 2011 |
| Ae-Ps7.6a | Baccara × PI 180693 and Baccara × 552 RIL | 233.9 | AB136 | 253,363,867 | SSR | 8.9 |  |
|  |  | 238.2 | AB122 | 261,521,510 | SSR | 3 |  |
| Ae-Ps7.6 | DSP x 90-2131 | 172.8 | AB27 | 286,432,964 | SSR | 10.7/5.9 | Hamon et. al., 2013 |

**Figure 8.**
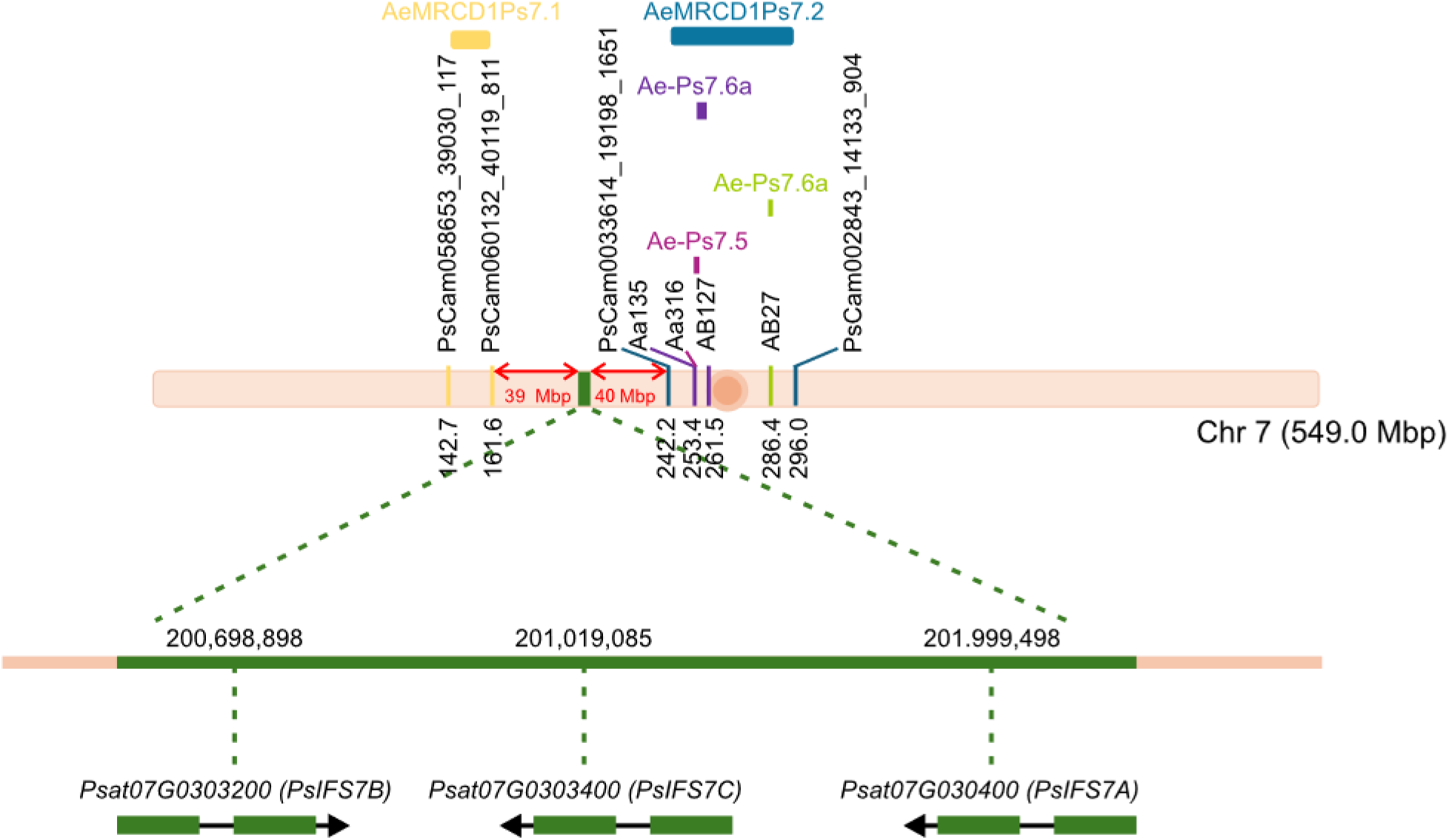
Genomic location of *PsIFS* genes and QTL associated with ARR resistance. Each QTL is color-coded based on its corresponding marker locations (in bp) on chromosome 7. The magnified region illustrates the precise genomic positions of three *PsIFS* genes (green) relative to neighboring markers.

## 4. Discussion

Isoflavonoids are well recognized for their health-promoting properties, including protection against cardiovascular diseases, hormone-dependent cancers, and osteoporosis in humans and animals (R. A. Dixon, 2004; R.A. Dixon & Ferreria, 2002; Messina, 1999). In plants, they function as signaling molecules in symbiotic interactions with nitrogen-fixing bacteria and exhibit antimicrobial activity that contribute to defense against a wide range of biotic stressors (Misra et al., 2010; Phillip, 1992; Samac et al., 2007; Subramanian et al., 2007). IFS, a P450 of the CYP93C subfamily, catalyzes the key branch-point reaction that diverts flavanones from the general flavonoid pathway toward isoflavonoid biosynthesis, leading to the production of the isoflavone aglycones daidzein and genistein.

Although *IFS* genes have been identified in several plant species, only a few of them have been functionally characterized (Sohn et al., 2021). Our genome-wide analysis identified three *IFS* genes in pea genome. Similarly, three *IFS* genes have been reported in *Vigna radiata*, *Phaseolus vulgaris*, *L. japonicus, Medicago sativa,* and *M. truncatula* while only two *IFS*s have been found in *G. max*, *G. soja*, and *L. culinaris* (Du et al., 2016; Jung et al., 2000; Shimada et al., 2000). All three *PsIFS* candidates are located on chromosome 7. Comparative sequence analysis revealed notable varietal polymorphisms in *PsIFS7A* and *PsIFS7B* between the reference genomes of pea cultivars Caméor and Zhongwan6. A previously described CYP93C18 from pea exhibiting IFS activity (Pičmanová et al., 2013) shares 98.1% and 97.9% nucleotide sequence identity with *PsIFS7B* and *PsIFS7C*, respectively and 94% with *PsIFS7A*. Because *CYP93C18* could not be uniquely mapped in the pea genome, it is plausible that *PsIFS7B* or *PsIFS7C* correspond to this previously reported gene.

Despite the conservation of the canonical P450 motifs, several CYP93C-specific alterations were detected among the PsIFS proteins. The Gly residue at the third position of the PPGP motif was substituted by Ser or Cys, consistent with similar substitutions reported in soybean CYP93 members (Khatri et al., 2022). Within P450 enzymes, the spacing between the oxygen-binding motif and the two K-helix motifs typically remains conserved within a subfamily (Hasemann et al., 1995; Khatri et al., 2022). A precise spacing between two consecutive NPA domains of aquaporins is crucial for their functionality (Deshmukh et al., 2015). In pea, all three PsIFS proteins maintained consistent spacing between two motifs except PsIFS7A, which contained an additional Gly residue (Gly424) between the fourth and fifth motifs, suggesting possible implications for catalytic efficiency or substrate orientation (**Figure 3, Supplementary Table S2**).

The tissue-specific expression analysis further supports potential functional differentiation among PsIFS isoforms. *PsIFS7A, PsIFS7B*, and *PsIFS7C* transcripts were most abundant in roots, with *PsIFS7A* also expressed in immature seeds (**Figure 5a**). This pattern of differential expression resembles that observed in soybean, where *GmIFS1* and *GmIFS2*, despite the fact that they share 93% nucleotide identity, display distinct tissue localization and catalytic properties. *GmIFS1* is primarily expressed in roots, whereas *GmIFS2* predominates in embryos and pods (Dhaubhadel et al., 2003; Subramanian et al., 2004; Subramanian et al., 2006). Furthermore, several *PsIFS* genes are highly induced upon pathogen challenge, consistent with their proposed role as entry points into pisatin biosynthesis. These results support the hypothesis that IFS activity represents a critical regulatory step in phytoalexin production, enabling rapid plant defense. Moreover, the differential expression patterns of PsIFS paralogs suggest potential sub-functionalization, which may fine-tune the biosynthetic response under specific biotic or environmental conditions.

Homology modeling highlighted differences in hydrophobic and aromatic interactions among PsIFS7A, PsIFS7B, and PsIFS7C, underscoring structural diversity within the PsIFS family. These results suggest that substrate recognition and stabilization may rely on residues adjacent to, rather than within, the canonical P450 motifs, representing structural adaptations that fine-tune flavanone specificity (Fürst et al., 2019; Khatri et al., 2023). Despite the high structural similarity among the predicted PsIFS proteins (**Figure 4**), PsIFS7A failed to catalyze the conversion of naringenin to genistein (**Figure 6**), indicating subtle but functionally significant alterations in its substrate-binding pocket that compromise catalytic activity.

Functional validation through *in vitro* enzymatic assays and overexpression of *PsIFS7C* in pea hairy roots confirmed the involvement of PsIFS enzymes in initiating the isoflavonoid biosynthetic pathway. Furthermore, the genomic proximity of *PsIFS* genes to QTLs associated with ARR resistance (**Figure 8**) provides additional evidence supporting their potential role in defense.

Collectively, these findings provide comprehensive insights into the molecular identity, structure-function relationships, and catalytic diversity of IFS isoforms in pea. The observed diversification within the pea IFS family may reflect adaptive evolution in response to distinct pathogen pressures or ecological niches. These insights can guide breeding strategies for disease-resistant pea varieties by targeting specific *PsIFS* alleles associated with enhanced pisatin accumulation.

## Acknowledgements

The authors thank Dr. Justin Renaud, Hodan Halane and Alex Molnar (AAFC-London) for technical assistance and Dr. Syama Chatterton (AAFC-Lethbridge) for pea germplasm. This research was funded by the Western Grains Research Foundation, Results Driven Agriculture Research, and Alberta Pulse Growers Commission through the Agriculture Funding Consortium (2022F108R) and AgriScience Program (J-003797) to SD.

## Data Availability Statement

The data that support the findings of this study are available from the corresponding author upon reasonable request.

## Supplementary Figures

**Supplementary Figure S1.**
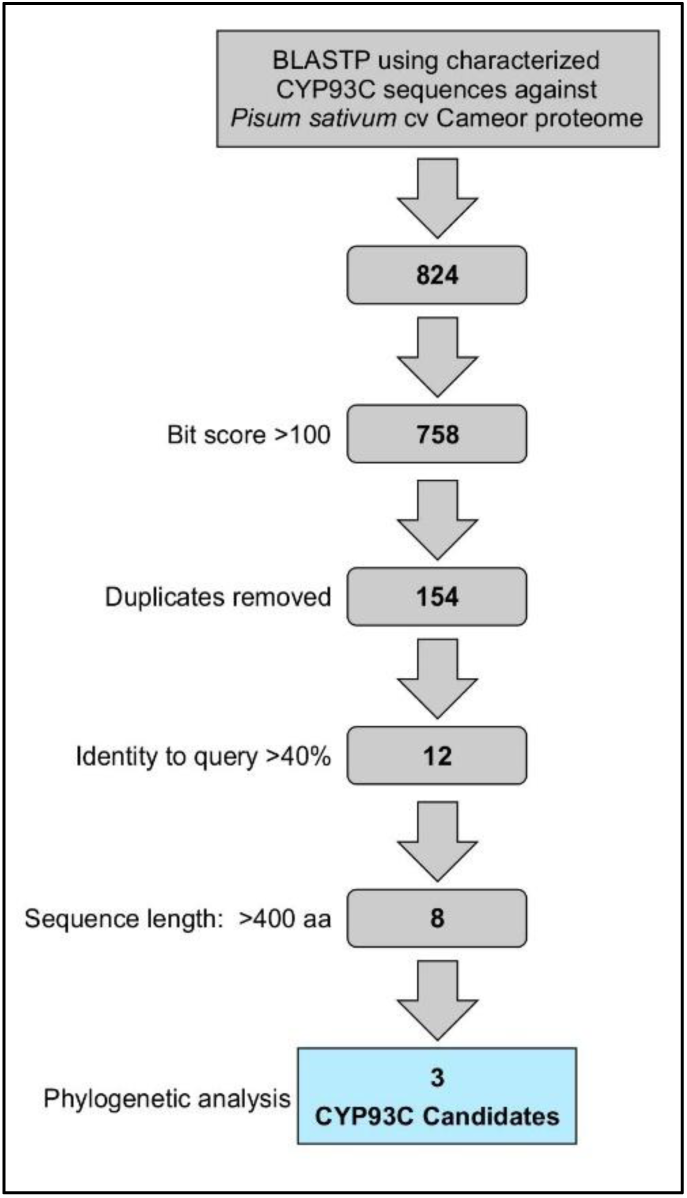
Strategy for identification of *PsCYP93C* (*PsIFS*) candidates in pea.

**Supplementary Figure S2.**
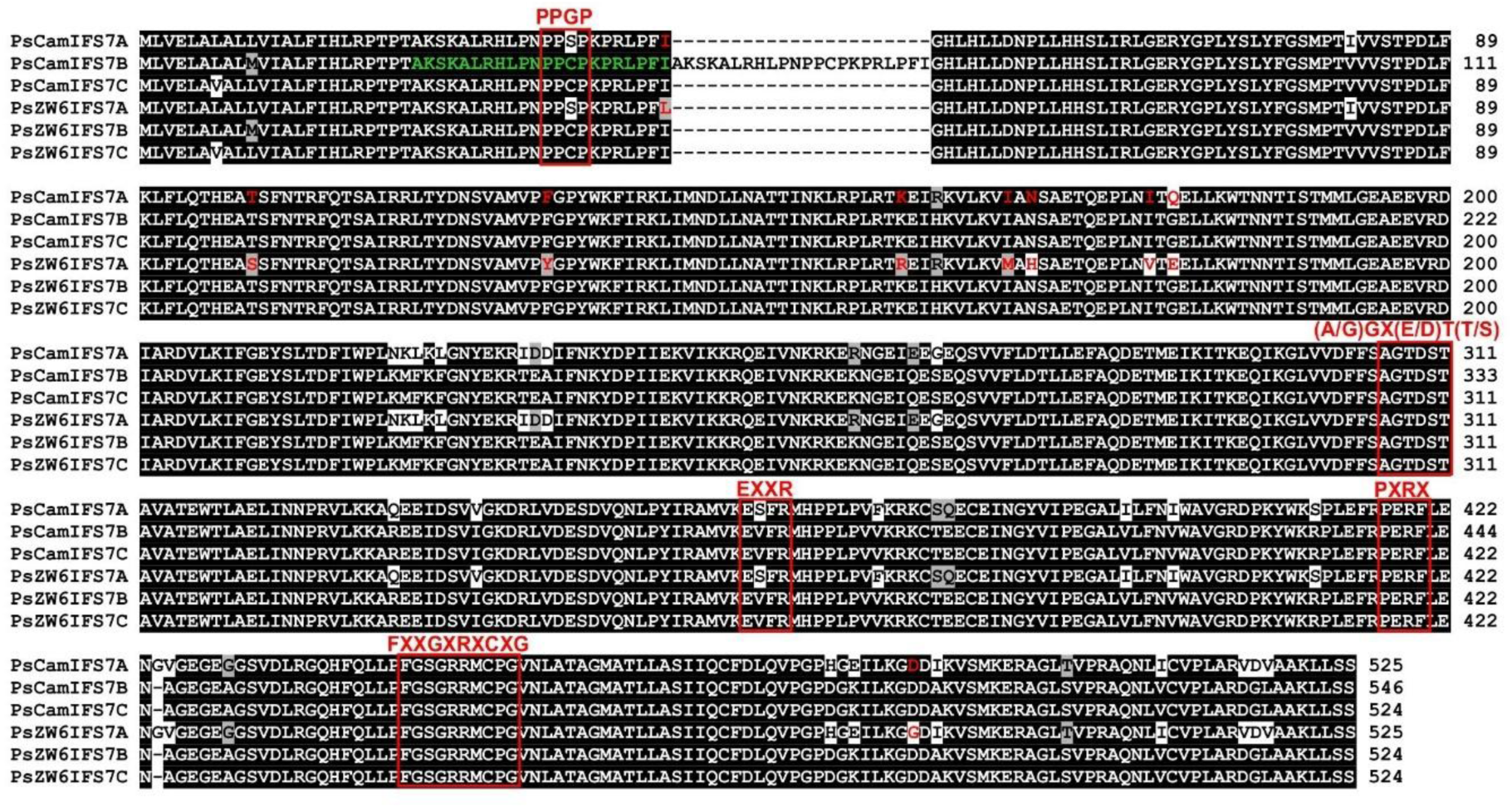
Sequence alignment of PsIFS proteins from Pea cv Caméor and Zhongwan6.

**Supplementary Table S1.** Identification of PsIFS candidates.

**Supplementary Table S2.** CYP93 protein sequences used for phylogenetic tree generation.

**Supplementary Table S3.** List of primer sequences.

